# A tale of two connectivities: intra- and inter-subject functional connectivity jointly enable better prediction of social abilities

**DOI:** 10.1101/2022.02.28.482399

**Authors:** Hua Xie, Elizabeth Redcay

## Abstract

Naturalistic functional magnetic resonance imaging (fMRI) paradigms, such as movie viewing, are attracting increased attention, given their ability to mimic the real-world cognitive demands on attention and multimodal sensory integration. Moreover, naturalistic paradigms allow for characterizing brain network responses associated with dynamic social cognition in a model-free manner using *inter-subject* functional connectivity (ISFC). While *intra-subject* functional connectivity (FC) characterizes the individual’s brain functional architecture, ISFC characterizes the neural coupling driven by time-locked extrinsic dynamic stimuli across individuals. Here, we hypothesized that ISFC and FC provide distinct and complementary information about individual differences in social cognition. To test this hypothesis, we examined a public movie-viewing fMRI dataset with 32 healthy adults and 90 typically developing children. Building three partial least squares regression (PLS) models to predict social abilities using FC and/or ISFC, we compared predictive performance to determine whether combining two connectivity measures could improve the prediction accuracy of individuals’ social-cognitive abilities measured by a Theory of Mind (ToM) assessment. Our results indicated that the joint model (ISFC+FC) yielded the highest predictive accuracy and significantly predicted individuals’ social cognitive abilities (rho = 0.34, *p* < 0.001). We also confirmed that the improved accuracy was not due to the increased feature dimensionality. In conclusion, we demonstrated that intra-/inter-subject connectivity encodes unique information about social abilities, and a joint investigation could help us gain a more complete understanding of the complex processes supporting social cognition.

## 1 Introduction

Humans are inherently social beings capable of drawing inferences about other people’s unobservable beliefs and intentions to navigate their social worlds. Using a suite of carefully designed cognitive tasks or resting-state paradigms, traditional functional magnetic resonance imaging (fMRI) studies have consistently identified a set of functional networks, termed the social brain, that supports this ability to perceive, understand, and react to the social world around us (Gallagher and Frith, 2003; Gobbini et al., 2007; Mitchell, 2008; Alcalá-López et al., 2018). While much of this work relies on artificial contexts, more recent novel paradigm-free functional neuroimaging such as naturalistic fMRI provides a mechanism to examine how the social brain may respond in complex, dynamic, real-world situations. With the dynamic and rich stimuli such as movies, stories, games, and virtual reality, naturalistic paradigms more closely mimic real-world demands on attention and multimodal sensory integration than the abstract and restricted stimuli employed in conventional fMRI paradigms, allowing for researchers to study humans’ highly interactive socio-cognitive processes “in the wild” (Finn et al., 2018, 2020; Razi et al., 2018; Richardson et al., 2018; Redcay and Moraczewski, 2020). In addition to the higher ecological validity, naturalistic fMRI also enables examining multiple neural measures (e.g., intra- and inter-subject neural coupling) to probe the mechanisms underlying the fleeting socio-cognitive information processing (Sonkusare et al., 2019). Despite the recent progress made, we argue that a gap of knowledge still exists since most of the previous naturalistic studies have looked at neural coupling measures in isolation (c.f., Simony et al., 2016; Kim et al., 2018; Lynch et al., 2018; Demirtaş et al., 2019), which begs the question of whether we can benefit from jointly studying the two neural coupling measures.

One such neural measure is intra-subject neural coupling, or functional connectivity (FC), computed as the temporal correlation of time-series across different brain regions within an individual. Intra-subject neural coupling is most commonly examined through resting-state FC (RSFC), which measures the intrinsic functional network architecture driven by spontaneous brain activities (Fox et al., 2005). Task-evoked FC, however, reflects brain network architecture patterns associated with a cognitive state, driven by a mixture of spontaneous and task-evoked brain activities (Cole et al., 2014). This task-evoked FC during movie-viewing, has shown great promises in studying individual differences as naturalistic paradigms may serve as an “amplifier” tapping into specific socio-cognitive domains, consistently outperforming RSFC (Vanderwal et al., 2017; Finn et al., 2020; Finn and Bandettini, 2021). For example, movie-viewing FC has led to more accurate predictions of phenotypes in the cognition and emotion domain (Finn and Bandettini, 2021).

Another neural measure is inter-subject neural coupling. Unique to naturalistic paradigms, inter-subject neural coupling characterizes brain responses associated with dynamic social cognition in a model-free manner (Nummenmaa et al., 2018), namely inter-subject correlation (ISC) and inter-subject functional connectivity (ISFC). Unlike the intra-subject FC, ISC and ISFC measure the neural coupling across individuals. ISC identifies the shared activation patterns of a given brain region across subjects (Hasson et al., 2004), and ISFC further delineates common functional connectivity patterns driven by the extrinsic time-locked dynamic stimuli (Simony et al., 2016). The shared activity and connectivity patterns across individuals reflect the shared understanding of the narratives (Nguyen et al., 2019), differ based on clinical diagnosis, e.g., autism (Salmi et al., 2013; Bolton et al., 2018) and attention-deficit/hyperactivity disorder (Salmi et al., 2020), is associated with collaboration outcomes (Xie et al., 2020), personality traits (Finn et al., 2018) and brain functional specialization in childhood (Moraczewski et al., 2018; Richardson, 2019).

Despite both intra- and inter-subject neural coupling showing great promise in furthering our understanding of individual differences in the neural underpinning of social cognition, only a handful of studies have systematically evaluated two types of neural coupling measures together (Simony et al., 2016; Kim et al., 2018; Lynch et al., 2018; Demirtaş et al., 2019). We argue a joint investigation of both neural coupling measures is needed because each measure may provide unique information about brain functions. Those studies that did examine both measures fell short of addressing the issue as they mainly focused on comparing ISFC and FC patterns. For instance, Lynch and colleagues found that ISFC patterns could not fully explain the FC changes during movie-viewing (Lynch et al., 2018). Similarly, Demirtaş and colleagues confirmed that while there was overlap between the two neural coupling patterns, increased coupling within frontal brain regions and reduced coupling between frontal-parietal brain regions were observed in intra-subject FC during movie-viewing (Demirtaş et al., 2019). Moreover, recent evidence has suggested that task-evoked FC modulation only accounts for a relatively small portion of individuals’ connectivity patterns during tasks, indicating that FC measured during tasks may still primarily reflect brains’ baseline functional architecture, i.e., FC fingerprint (Gratton et al., 2018; Xie et al., 2018). In another study comparing two neural patterns, Simony and colleagues demonstrated the greater sensitivity of ISFC than standard intra-subject FC directly measured during movie-viewing in detecting stimulus-induced connectivity patterns (Simony et al., 2016).

Taken together, while the two neural coupling measures are clearly distinct from one another in some ways, there is a gap in our knowledge concerning whether the information encoded in these two connectivity patterns is complementary or redundant. To answer our question, we used an open-access movie-viewing fMRI dataset and built a partial least square (PLS; Krishnan et al., 2011) regression model to predict children’s socio-cognitive abilities using intra- and/or inter-subject connectivity. Here, we focused on the social brain network (Alcalá-López et al., 2018), which undergoes rapid functional specialization throughout childhood while children gain social skills (Gweon et al., 2012; Gweon and Saxe, 2013; Richardson et al., 2018). We postulated that if both connectivity measures provided complementary information about individuals’ social abilities, the joint model with both connectivity measures would offer the highest predictive performance. We also conducted confirmation analysis to verify previous studies’ observations on the similarity between ISFC and FC patterns (Simony et al., 2016; Kim et al., 2018; Lynch et al., 2018; Demirtaş et al., 2019) and carried out additional exploratory analysis to assess the neurobiological significance of such similarity.

## 2 Materials and Methods

### 2.1 Participants

An open-access dataset was used in this study (https://www.openfmri.org/dataset/ds000228/), which contains a large sample of children (n = 122, 3.5 - 12 years old, 64 females, nine left-handed), and adults (n = 33, 18–39 years old, 20 females). No participants had any known cognitive or neural disorders. All adult participants and the parent/guardian of the child participants gave written consent. All protocols were approved by the Committee on the Use of Humans as Experimental Subjects at the Massachusetts Institute of Technology.

### 2.2 Movie task & behavioral battery

Participants underwent fMRI scans while watching a short silent version of “Partly Cloudy,” a 5.6-min animated movie with plots eliciting frequent inferences of characters’ mental states (beliefs, desires, emotions) and bodily sensations (particularly pain). The movie began after 10 seconds of a black screen and 10 seconds of opening credits. After fMRI scans, all children completed a socio-cognitive behavioral battery measuring theory of mind (ToM) abilities (Gweon et al., 2012). The custom-made ToM battery (available at https://osf.io/G5ZPV/) involved listening to an experimenter tell a story and answering prediction and explanation questions that required reasoning about characters’ mental states.

### 2.3 Image acquisition and preprocessing

Whole-brain structural and fMRI data were acquired on a 3-Tesla Siemens Tim Trio scanner at the Massachusetts Institute of Technology. All participants were scanned using the standard Siemens 32-channel head coil except for those under age five, who used custom 32-channel phased-array head coils made for younger children. T1-weighted structural images were collected in 176 interleaved sagittal slices with 1 mm isotropic voxels (Adults: FOV = 256 mm; children: FOV = 192 mm). Functional data were collected with a gradient-echo EPI sequence (#slices = 32; TR = 2 s, TE = 30 ms, flip angle = 90°). The details of scan protocols can be found in (Richardson et al., 2018).

We used the data preprocessed by the original authors (Richardson et al., 2018). Specifically, all functional images were first registered to the Montreal Neurological Institute (MNI) template, and registration of each individual’s brain to the MNI template was visually inspected. The registered data were then smoothed using a 5 mm Gaussian kernel. The Artifact Detection Tools (https://www.nitrc.org/projects/artifact_detect/) were used to detect timepoints with more than 2 mm framewise displacement (FD) to the previous time point or with a fluctuation in the global signal that exceeded a threshold of three standard deviations from the mean global signal. Additionally, we applied temporal interpolation on artifactual timepoints and regressed out the first five principal components of the white matter signal from the white matter signal CompCor (Behzadi et al., 2007). The residual time courses were then band-pass filtered between 0.008-0.15Hz and detrended with the first- and second-order polynomials. We also excluded the volumes corresponding to the opening credits (first 10 TRs). We adopted a stringent exclusion criterion and removed participants with a mean FD greater than 0.5mm, leaving 90 child and 32 adult participants for further analysis. We performed scrubbing by removing TRs with FD greater than 1mm.

### 2.4 Intra- and inter-subject functional connectivity

Following preprocessing, we extracted the ROI timeseries from the denoised data using the social brain atlas (Alcalá-López et al., 2018) containing 36 regions of interest (ROIs). The social brain atlas, developed using meta-analyses of social and affective abilities, consists of key regions responsible for social cognition, such as the amygdala, precuneus, MPFC, and TPJ, among many other regions responsible for social and affective information processing. The ROI masks were dilated by one voxel along each direction, and then masked by a binary group mask, resulting in 4 ROIs (bilateral temporal pole and cerebellum) being dropped due to little spatial coverage (fewer than five voxels). Next, we extracted the mean timeseries from each ROI mask. As shown in Fig.1, we computed the intra-subject FC by correlating the timeseries using Pearson correlation. For the inter-subject connectivity, Pearson correlation was computed across different individuals (i.e., child-to-adult) within a given ROI (ISC) and across two ROIs (ISFC). More specifically, we used the adults’ timeseries as a reference, and we correlated the children’s timeseries with those of the adults. We chose adults as the reference group, given the previous evidence showing stronger and more coherent neural coupling and activation patterns in adults than children (Cantlon and Li, 2013; Moraczewski et al., 2018). We averaged upper- and lower-diagonal ISFC values to obtain symmetric ISFC matrices. Both connectivity measures were then Fisher Transformed and normalized to have zero mean and unit variance. Since ISC corresponds to the diagonal terms of the ISFC matrix, hereinafter, we will use ISFC to refer to both ISC and ISFC.

**Figure 1.**
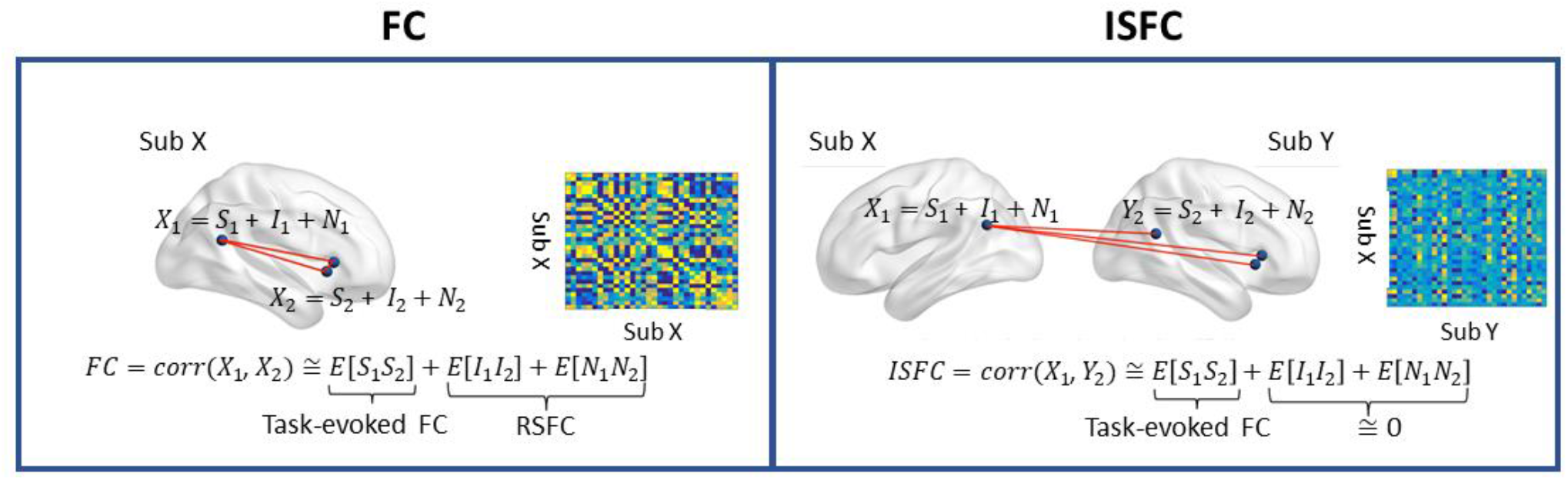
A graphical depiction of intra-subject functional connectivity (FC) and inter-subject ISFC between two ROIs. *S:* task-evoked brain activity; *I:* intrinsic brain activity; *N:* noise. Left: FC is characterized as the neural coupling within a subject, and the RSFC is measured in the absence of the task-evoked signal (*S*). Right: ISFC measures the brain synchronization across subjects, as only the neural coupling driven by the task-evoked brain activity is preserved, as the intrinsic brain activity and noise are uncorrelated across individuals. The ISC corresponds to the diagonal terms of ISFC. Figure adapted from (Simony et al., 2016).

### 2.5 Building prediction model using partial least square regression

PLS regression is a statistical method that identifies a linear relationship between predictors (*X*, connectivity) and response (*Y*, ToM scores). Unlike principal component analysis (PCA), which finds the latent factors that explain the most variance in *X*, PLS regression identifies the latent components that best associate predictors with the response. For a univariate response, PLS regression finds an orthonormal combination of *X* termed as *w*, which maximizes the covariance matrix of *cov*(*Xw,Y*).

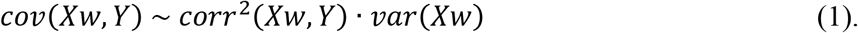

The *w* in Eq (1) can be solved using singular value decomposition.

PLS is well-suited for questions where the dimensionality of independent variables (#connectivity) is much higher than the number of observations (#participants), and has been previously used in many associating brain patterns with behavior (Yoo et al., 2017; Ching Fong et al., 2018).

Here, we built three models, namely FC, ISFC, the joint model (ISFC+FC), to predict the ToM scores after controlling for age (linear and quadratic), gender, and handedness. Ten-fold cross-validation was used to determine the optimal dimensionality of the latent components, and the cross-validation was repeated 100 times to minimize the influence of sampling variability. The model performance was evaluated by the mean absolute error (MAE) and Spearman correlation between the true and predicted ToM scores. To evaluate prediction performance for the chosen analyses, we ran 50,000 permutation tests to derive empirical null distributions of ToM prediction. We also conducted a partial correlation analysis to ensure that age and head motion did not drive our prediction.

## 3 Results

### 3.1 The joint model best predicted ToM scores

We compared the predictive performance of three models across a wide range of hyperparameters (#components = 1-10). As shown in Fig. 2(a&b), the optimal predictive performance, as reflected by the lowest mean absolute error (MAE) and highest correlation between the true and predicted ToM scores, was achieved with the joint model (ISFC+FC) with the component number equal to 2. Permutation results revealed that the optimal model significantly predicted ToM scores (mean rho = 0.34, *p* < 0.001), as shown in Fig. 2c.

**Figure 2.**
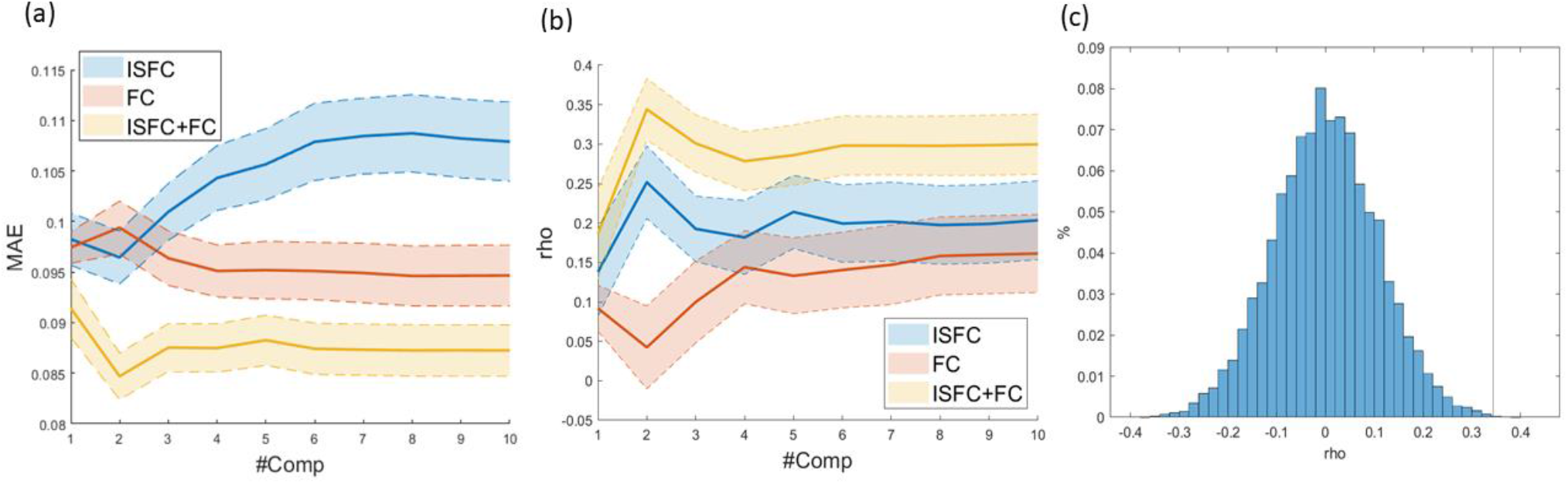
(a) Average mean absolute error (MAE) as a function of the number of latent components (#comp) over 100 10-fold cross-validations. The shaded area indicates one standard deviation. (b) Spearman correlation between true and predicted ToM scores. (c) Null distributions of ToM scores with the vertical line indicating actual model performance.

To rule out the possibility that the increased feature dimensionality drove the improved predictive performance, we randomly sampled half of the FC and ISFC edges and re-evaluated the model performance 100 times. As shown in Fig. 3, despite the slight drop in predictive performance, the joint model using only half of the FC and ISFC features still outperformed the ISFC and FC model separately (mean rho = 0.32).

**Figure 3.**
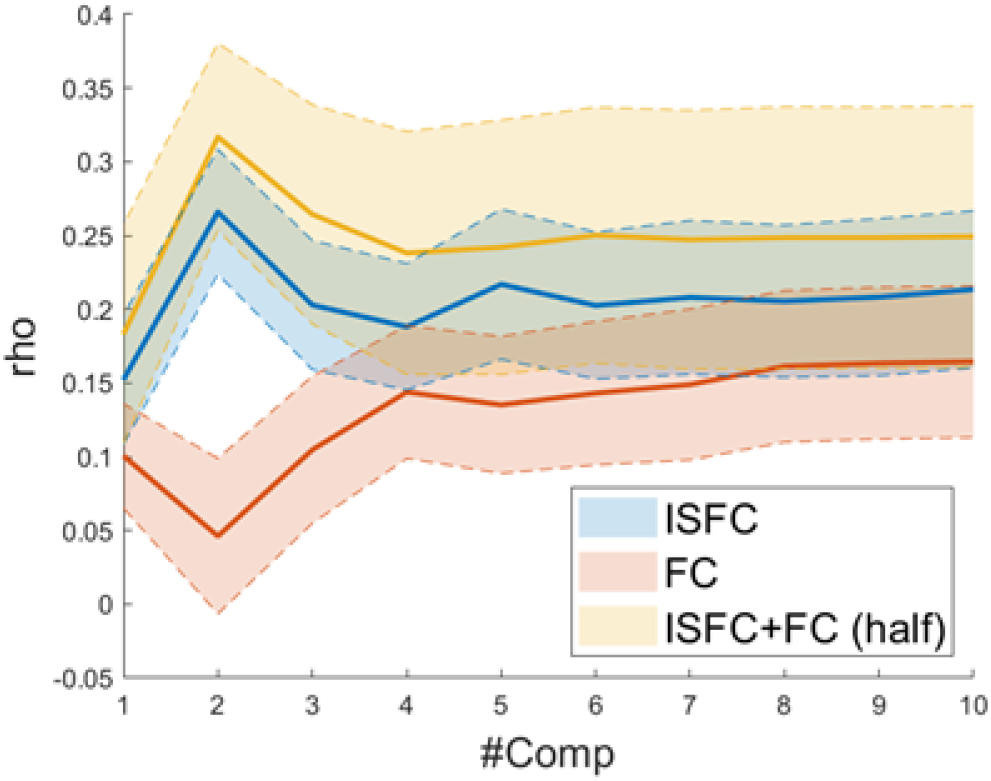
The joint model using only half of the features still outperformed the FC and ISFC model (paired t-test *ps* < 0.001), suggesting that the better predictive performance was not merely caused by the increased number of features of the joint model.

**Figure 4.**
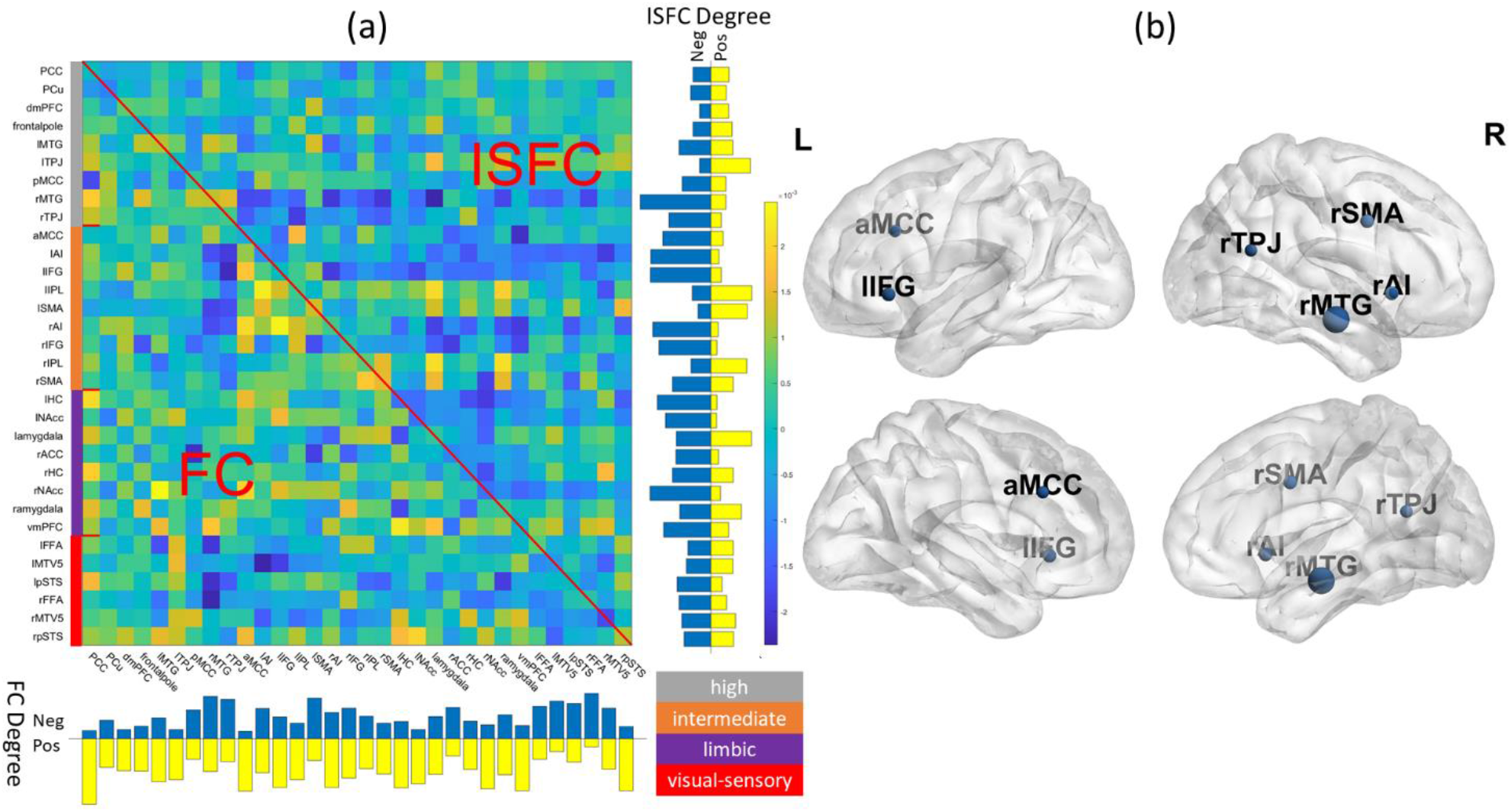
(a) Visualization of beta coefficients of the final joint model and the degree (row sum/column sum) of ISFC and FC patterns. The network assignment was color-coded based on the definition of (Alcalá-López et al., 2018). Grey: high-level processing; orange: intermediate-level processing; purple: limbic; red: visual-sensory. (b) Key regions identified in the final model (Bonferroni corrected *p* < 0.05), and the node size is proportional to the drop in predictive accuracy. For ROI abbreviations, see Table S1.

### 3.2 Leave-one-region out analysis identifying key regions in the predictive model

We averaged 100 joint models with two components and visualized associated beta coefficients as shown in Fig. 3a and showed identified latent components in Fig. S1. As is typical of a data-driven approach, regions with high beta values were diffusive across the social brain, with no single dominant anatomical pattern. To better delineate the key ROIs that contributed to the prediction accuracy, we conducted a leave-one-out (LOO) analysis by removing each ROI at a time and compared the predictive performance of the reduced model against the full joint model. We found that excluding anterior midcingulate cortex (aMCC), right middle temporal gyrus (rMTG), right TPJ (rTPJ), right anterior insula (rAI) and right supplementary motor area (rSMA), and left inferior frontal gyrus (lIFG) significantly lowered the predictive performance (Bonferroni corrected *p* < 0.05), suggesting the significance of these regions in our predictive model. We also conducted a follow-up LOO analysis excluding ISFC and FC edges separately. We found that excluding FC profiles of rMTG and lIFG as well as the ISFC profiles of rMTG, rSMA, and ventromedial prefrontal cortex (vMPFC) significantly lowered the model’s prediction performance (Bonferroni corrected *p* < 0.05).

### 3.3 On FC-ISFC similarity

Given the previous evidence showing high similarity between ISFC (excluding diagonal ISC) and intra-subject movie-FC patterns (Kim et al., 2018; Demirtaş et al., 2019), we conducted a similar confirmation analysis by comparing the two connectivity patterns, which revealed a moderate positive correlation (mean rho = 0.34). Moreover, it should be noted that the FC-ISFC similarity is modulated by the diagonal ISC terms (for a detailed explanation, see supplemental materials), and higher ISC translates to more adult-like brain responses in children. We were interested in examining whether ISFC-FC similarity had any neurobiological significance, given the previous studies showing children with better task performance had more adult-like brain responses (Cantlon and Li, 2013; Cai et al., 2019). Thus, we carried out a follow-up exploratory analysis by correlating individuals’ ISFC-FC similarity with age or ToM abilities. We observed the similarity between ISFC and FC to be positively correlated with age (rho = 0.550, *p* < 0.001), while not correlated with ToM scores (rho = 0.12, *p* > 0.05).

## 4 Discussion

Previous research on the neural underpinnings of socio-cognitive information processing has typically relied on carefully constructed cognitive tasks and task-free paradigms (resting-state). However, these approaches are limited in that they have poor ecological validity. The emergence of naturalistic fMRI paradigms offers many theoretical and practical advantages, one of which is simultaneous examining intra-&inter-subject connectivity, namely FC and ISFC. However, no studies so far have examined the potential synergy of combining these two connectivity measures and whether they encode complementary information. Here, using the predictive performance as a proxy of features’ informativeness, we filled the knowledge gap by comparing the predictive efficacy of models using features of FC and/or ISFC to predict children’s social-cognitive abilities (ToM scores) using a social brain atlas (Alcalá-López et al., 2018). Moreover, we also delineated key ROIs in our predictive model to better understand regions’ contribution to individual differences in social abilities.

### A synergy of intra-&inter-subject FC

By comparing the predictive efficacy of FC and ISFC, our results suggested that there may indeed be synergy between the ISFC and FC to predict individual differences. As shown in Fig. 2, the joint model using ISFC and FC not only significantly predicted participants’ ToM scores, but far exceeded individual models’ predictive accuracy. More importantly, the reduced joint model, which roughly matched the dimensionality of ISFC and FC, still outperformed individual models by a wider margin, as shown in Fig. 3 (*ps* < 0.001). We speculate that the superior performance of the model combining the connectivity measures may indicate that distinct cognitive factors may drive these two neural couplings. ISFC theoretically more precisely captures the shared connectivity patterns driven by the dynamic movie stimuli than standard task-FC (Simony et al., 2016), since the physiological and motion artifacts were uncorrelated across subjects. Moreover, empirical evidence has shown that the time-varying ISFC better tracked the movie cues than the time-varying FC (Bolton et al., 2020), and ISFC within DMN exhibited reliable and distinct patterns during narrative processing (Simony et al., 2016), suggesting that ISFC may be a more cognitively relevant representation of movie-evoked connectivity pattern than the direct measurement of movie-FC. On the other hand, the intra-subject FC measured during movie-viewing is a combination of task-evoked FC and individual FC fingerprint, with the FC fingerprint being the dominant one (Gratton et al., 2018). Thus, including FC measures in our model provides additional information about individuals’ FC fingerprints to facilitate the prediction. As a previous study has shown the improved performance of combining intra-subject FCs from multiple conditions for predictive modeling of phenotypic measures (Gao et al., 2019), we further demonstrated the synergy of intra- and inter-subject FC for predicting socio-cognitive abilities using a movie-watching naturalistic paradigm.

To further elucidate the relationship between the two neural coupling measures, we examined the similarity of individuals’ ISFC (i.e., excluding the diagonal ISC terms) and FC patterns, which previous studies have focused on (Kim et al., 2018; Lynch et al., 2018; Demirtaş et al., 2019). Consistent with the prior work, we found that the two connectivity patterns were positively correlated (Kim et al., 2018; Demirtaş et al., 2019, c.f., Lynch et al., 2018). Both ISFC and FC patterns contained some degree of movie-evoked connectivity, which could have given rise to the overall positive ISFC-FC similarity. We also found that FC-ISFC similarity was significantly positively correlated with children’s age, suggesting more adult-like social brain responses in older children, consistent with the earlier observation (Richardson et al., 2018). Moreover, younger children could have less stable movie-evoked FC patterns as the underlying cognitive processes may be more variable and individualized, while stronger and more coherent movie-evoked FC patterns emerged in the older children, leading to a higher FC-ISFC similarity (Cantlon and Li, 2013; Moraczewski et al., 2018). Nevertheless, we noted that although FC-ISFC similarity was significantly correlated with age, there was no significant correlation between FC-ISFC and ToM scores, justifying the necessity of a more complex predictive model.

### Highly predictive nodes concentrated in intermediate and high-level processing subnetworks

Our LOO analysis identified a few critical ROIs that significantly lowered the predictive accuracy in our final joint model. The majority of these key ROIs were found to be part of intermediate (i.e., rAI, aMCC, rSMA, and lIFG) and high-level processing subnetworks (i.e., rTPJ and rMTG), as defined by Alcalá-López et al. (2018). The high-level processing subnetwork consists of key regions closely associated with ToM, such as TPJ, MTG, posterior cingulate cortex (PCC), and precuneus (Schurz et al., 2014). Per Alcalá-López and colleagues, this high-level subnetwork is more strongly connected within itself than any other subnetworks, and is most likely to be associated with the cognitive categories of social cognition in all subnetworks (Alcalá-López et al., 2018). The significance of the high-level processing network was confirmed when examining the latent components in the final model, which associates the connectivity profiles to ToM scores, as the second component was weighted heavily towards ROIs within the high-level processing subnetwork (Fig. S1). Our LOO analysis also identified a few nodes from an intermediate-level subnetwork. The intermediate-level subnetwork, including AI, aMCC, IFG, among others, intertwines with lower-level visual-sensory subnetwork that handles the preprocessed social-affective environmental inputs (Alcalá-López et al., 2018). AI is known for its role as a bridge between large-scale brain networks. Together with aMCC, these two ROIs within the intermediate networks were involved in empathy and pain-related processing (Kurth et al., 2010; Lamm et al., 2011), and were also found among the ROIs that were highly predictive of ToM scores. Moreover, the absence of lower-level visual-sensory ROIs among the most predictive regions suggests that although movie stimuli induce wide-spread connectivity changes in visual areas within and across individuals (Hasson et al., 2004; Lynch et al., 2018), such connectivity changes may be less socio-behaviorally relevant as compared to those in the high association cortices.

### Limitations and future directions

Several limitations require further consideration. Firstly, we used adults as the reference group when computing ISFC. Our choice was justified by previous studies showing children with more variable inter-subject neural coupling and weaker activation patterns than adults across many cortical regions (Cantlon and Li, 2013; Moraczewski et al., 2018). Therefore, we believe the adults may be well-suited as a reference group given the higher homogeneity. Future investigation could focus on the impact of choosing different reference groups and better delineating the developmental effect. A second limitation is that we chose a set of a-priori ROIs using the social brain atlas since we are interested in predicting individuals’ social abilities. Future studies could examine a different parcellation, such as whole-brain parcellations (Power et al., 2011; Craddock et al., 2012; Shen et al., 2013), given the success of whole-brain FC predictive models (Shen et al., 2017; Beaty et al., 2018; Lake et al., 2019; Xiao et al., 2021). Alternatively, future studies could use atlases developed specifically for children since the brain parcellation used in the current study was derived from adult studies or conduct hyperalignment for better functional alignment (Haxby et al., 2020). Thirdly, we used PLS regression to extract the brain-behavior relationship while alternative learning models may improve predictive performance, such as sparse group LASSO (Bai et al., 2020) and sparse tensor decomposition (Zhang et al., 2021). Lastly, since the primary purpose of this study is to investigate the efficacy of combining two connectivity measures to predict social abilities, we did not include an external validation set to examine the robustness of our predictive model. Future studies with an independent validation set or different movie clips would further ensure the reproducibility of our findings.

### Conclusion

The current study investigated the potential benefits of jointly studying intra-&inter-subject connectivity, namely, FC and ISFC, respectively, and whether the two connectivity measures combined could enhance the prediction of individuals’ socio-cognitive abilities as measured by ToM scores. Using PLS regression, we showed that the joint connectivity (ISFC+FC) model outperformed individual models even after matching the feature dimensionality. Our results suggest that intra-&inter-subject connectivity may encode unique and complementary information about the individuals’ social abilities, and we shall make full use of both connectivity measures to gain enriched insight into neural processes underlying naturalistic fMRI paradigms.

## 5 Funding

Research reported in this publication was supported by the National Institute Of Mental Health of the National Institutes of Health under Award Number R01MH107441 and R01MH125370. The content is solely the responsibility of the authors and does not necessarily represent the official views of the National Institutes of Health.

## Data Availability Statement

The datasets analyzed for this study can be found on the OpenfMRI repository at https://www.openfmri.org/dataset/ds000228/.

---

**Figure S1.**
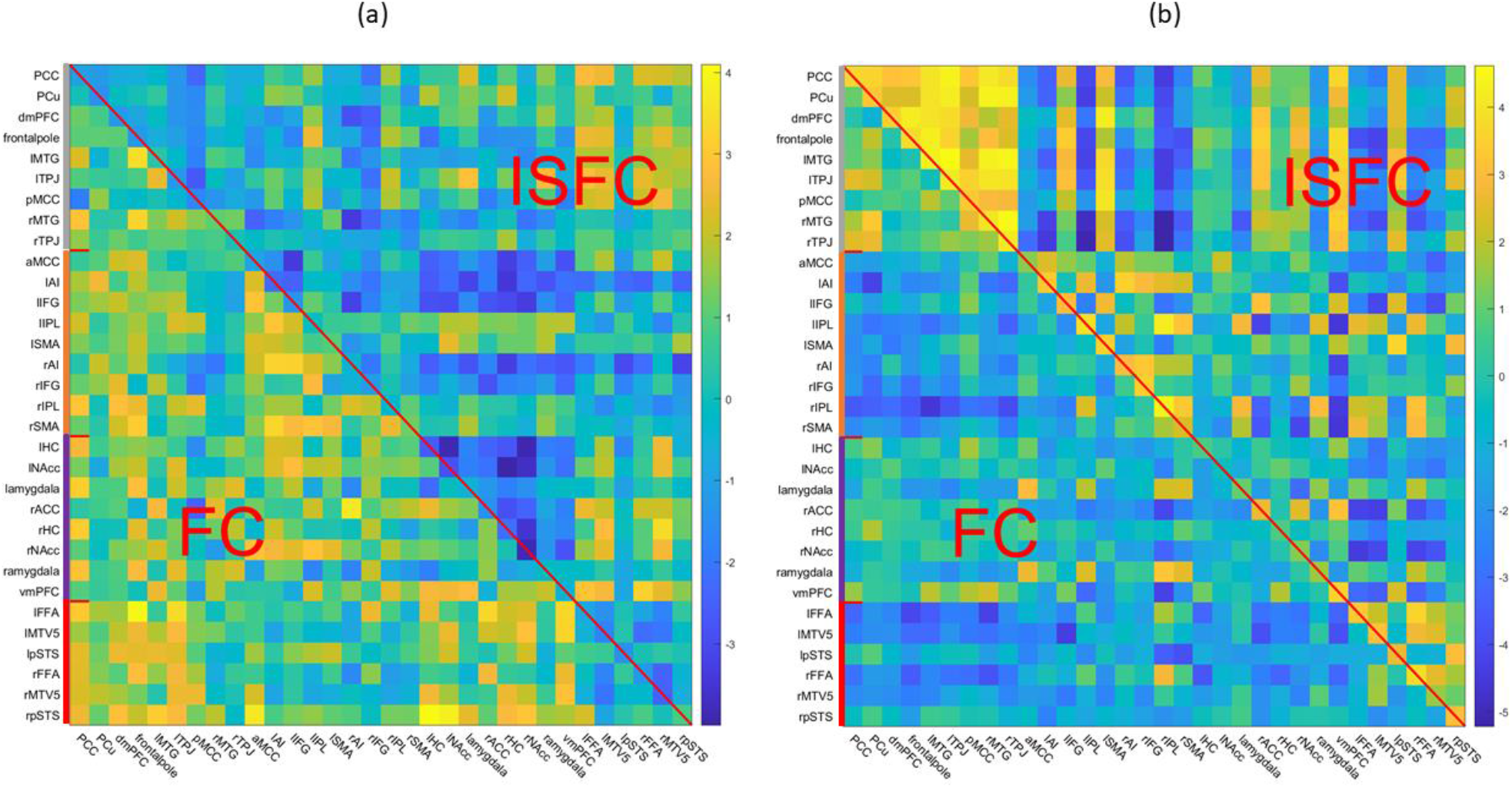
The two latent components identified in the final joint model. (a) The first latent factor explained 2.75% of the total connectivity variance, which mostly loaded onto the intra-subject neural coupling (FC). (b) The second latent factor explained 5.06% of the total connectivity variance, which loaded onto the inter-subject neural coupling (ISFC). The network assignment was color-coded based on the definition of (Alcalá-López et al., 2018). Grey: high-level processing; orange: intermediate-level processing; purple: limbic; red: visual-sensory.

**Table S1.**
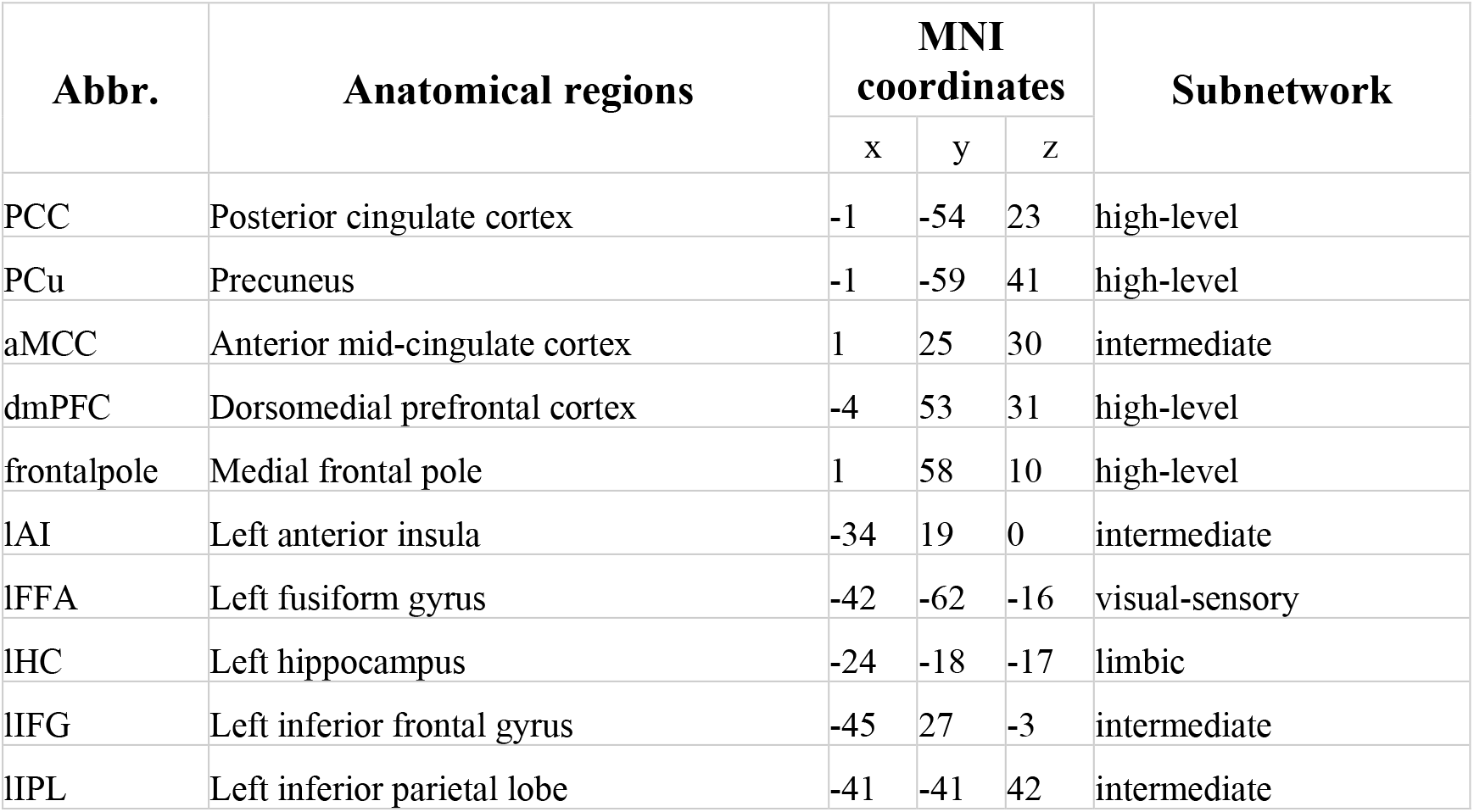
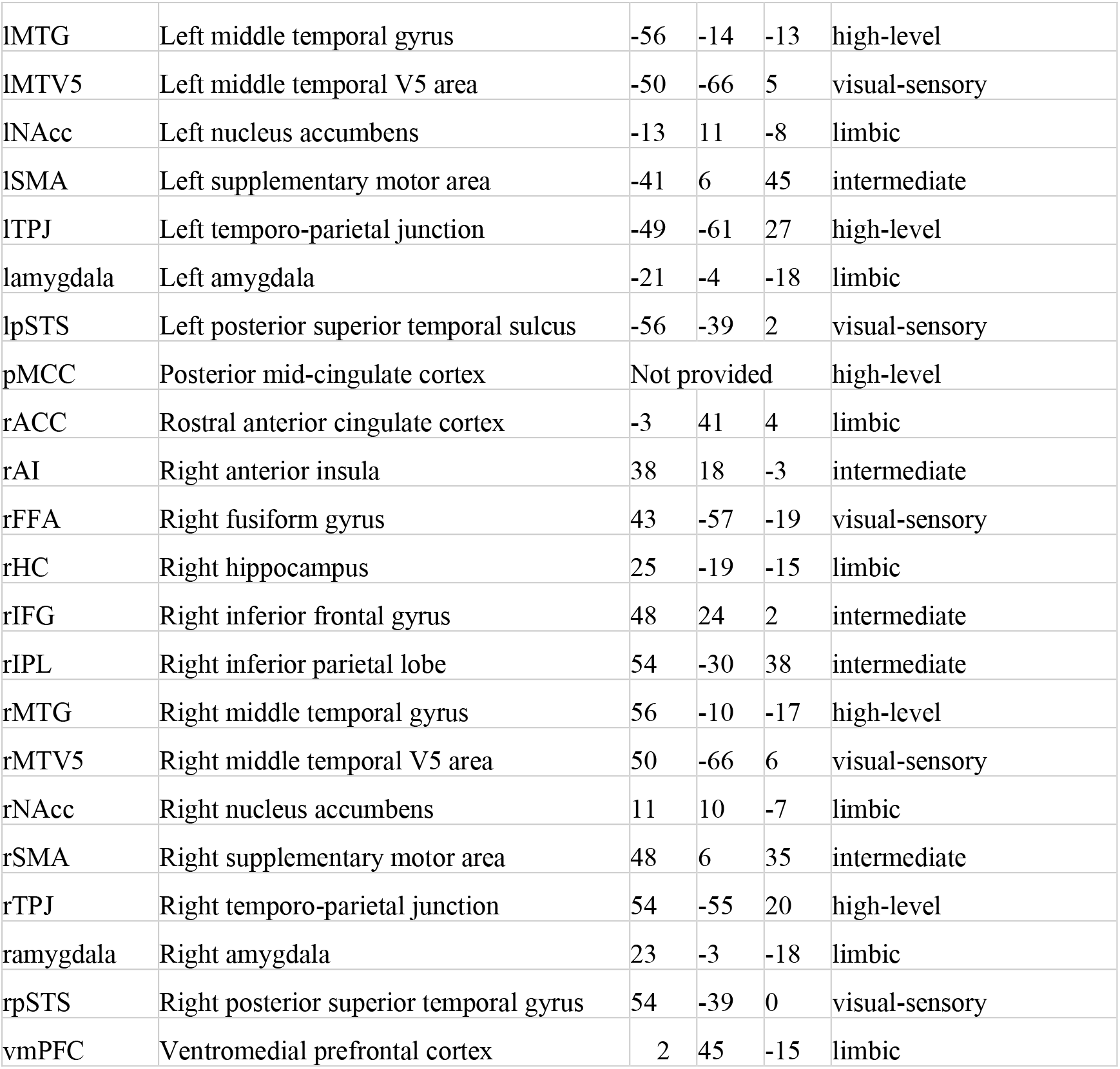
The thirty-two ROIs used in the study from the social brain atlas from (Alcalá-López et al., 2018). The parcellation is publicly available from the data-sharing platforms ANIMA (http://anima.fz-juelich.de/) and NeuroVault (http://neurovault.org/collections/2462/).

## Supplemental materials

On the relationship between FC, ISC and ISFC. By definition, we have

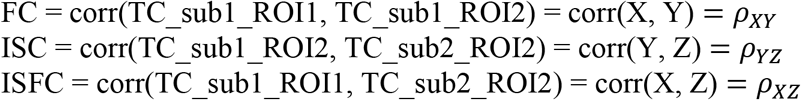

For simplicity, the corresponding time courses are referred to as X, Y and Z, and have zero mean and unit variance. Suppose we know FC (*ρ_XY_*) and ISC(*ρ_YZ_*), we can then express X and Z as follows

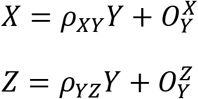

where 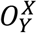 is the component of X that is orthogonal to Y (error). Then we can write ISFC (*ρ_XY_*) as

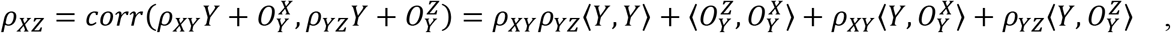

where 〈*x, y*〉 is the dot product of the two variables.

Recall that all three variables are of zero mean and unit variance, and both 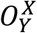 and 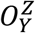 are orthogonal to Y, thus we can simplify *ρ_XZ_* as

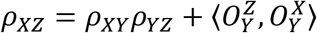

Using Cauchy-Schwarz inequality we can show that

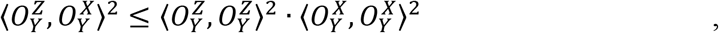

where 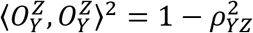, and 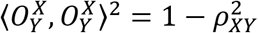.

Thus, we can show that ISFC (*ρ_XZ_*) is within a certain range depending on FC and ISC.

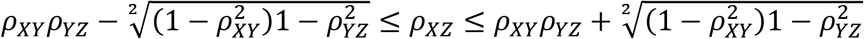

## Notes

### Competing Interest Statement

The authors have declared no competing interest.

